# Causal Association between Birth Weight and Adult Diseases: Evidence from a Mendelian Randomisation Analysis

**DOI:** 10.1101/447573

**Authors:** Ping Zeng, Xiang Zhou

## Abstract

**Background:** It has long been hypothesized that birth weight has a profound long-term impact on individual predisposition to various diseases at adulthood: a hypothesis commonly referred to as the fetal origins of adult diseases. However, it is not fully clear to what extent the fetal origins of adult diseases hypothesis holds and it is also not completely known what types of adult diseases are causally affected by birth weight. Determining the causal impact of birth weight on various adult diseases through traditional randomised intervention studies is a challenging task.

**Methods:** Mendelian randomisation was employed and multiple genetic variants associated with birth weight were used as instruments to explore the relationship between 21 adult diseases and 38 other complex traits from 37 large-scale genome-wide association studies up to 340,000 individuals of European ancestry. Causal effects of birth weight were estimated using inverse-variance weighted methods. The identified causal relationships between birth weight and adult diseases were further validated through extensive sensitivity analyses and simulations.

**Results:** Among the 21 adult diseases, three were identified to be inversely causally affected by birth weight with a statistical significance level passing the Bonferroni corrected significance threshold. The measurement unit of birth weight was defined as its standard deviation (i.e. 488 grams), and one unit lower birth weight was causally related to an increased risk of coronary artery disease (CAD), myocardial infarction (MI), type 2 diabetes (T2D) and BMI-adjusted T2D, with the estimated odds ratios of 1.34 [95% confidence interval (CI) 1.17 - 1.53, *p* = 1.54E-5], 1.30 (95% CI 1.13 - 1.51, *p* = 3.31E-4), 1.41 (95% CI 1.15 - 1.73, *p* = 1.11E-3) and 1.54 (95% CI 1.25 - 1.89, *p* = 6.07E-5), respectively. All these identified causal associations were robust across various sensitivity analyses that guard against various confounding due to pleiotropy or maternal effects as well as inverse causation. In addition, analysis on 38 additional complex traits found that the inverse causal association between birth weight and CAD/MI/T2D was not likely to be mediated by other risk factors such as blood-pressure related traits and adult weight.

**Conclusions:** The results suggest that lower birth weight is causally associated with an increased risk of CAD, MI and T2D in later life, supporting the fetal origins of adult diseases hypothesis.

## Background

Birth weight is a widely used surrogate measurement of intrauterine exposure and early life development, which has long been hypothesized to have a profound long-term impact on individual’s predisposition to various diseases at adulthood — a hypothesis commonly referred to as the fetal origins of adult diseases [1-4]. Indeed, early registry and other observational studies have provided strong empirical evidence supporting an inverse association between birth weight and the risks of several adult diseases [2-5]. Exemplary birth weight associated adult diseases include lung disease [6], coronary artery disease (CAD) and stroke [7-10], blood pressure [11, 12], type 2 diabetes (T2D) [13, 14] and asthma [15, 16]. However, it is unclear whether the identified associations between birth weight and the aforementioned adult diseases represent truly causal relationship, or are merely spurious associations caused by common confounding factors that occur during prenatal or postnatal life [3, 17-20] or confounding due to pleiotropy and shared genetic components [21]. Common confounding factors such as early life environment, lifestyle, current body mass index (BMI), or adult weight can be associated with both birth weight and adult diseases to cause spurious association between the later two [22]; and these confounding factors are often difficult to fully control for in observational studies [2]. As a consequence, some identified associations between birth weight and adult diseases in early studies have not been validated in recent studies. For example, the inverse association between birth weight and adult blood pressure identified in early studies are later found to be a consequence of failure to adjust for adult weight or other confounders [2,20, 23]. As another example, potentially due to different confounding effects, different studies show conflicting results with regard to the association between birth weight and T2D: T2D risk is positively associated with birth weight in some studies but negatively associated with birth weight in others [24, 25]. Therefore, it is not fully clear to what extent the fetal origins of adult diseases hypothesis holds and it is also not completely clear what types of adult diseases are causally affected by birth weight [17, 18].

Understanding the long-term causal impact of birth weight on individual’s predisposition to various disease risks is important from a public health perspective; as a better understanding can pave ways for using early nutritional intervention that can potentially increase birth weight to reduce disease burden in later life [26]. However, determining the causal impact of birth weight on various adult diseases through traditional randomised intervention studies is challenging, as such studies necessarily require long-term follow-ups, are time-consuming, expensive, and often times, unethical [21, 27, 28]. Therefore, it is desirable to determine the causal relationship between birth weight and various adult diseases through observational studies [29]. A powerful statistical tool to determine causal relationship and estimate causal effects in observational studies is Mendelian randomisation (MR). MR adapts the commonly used instrumental variable analysis method developed in the field of causal inference to settings where genetic variants are served as instrumental variables [30, 31]. In particular, MR employs genetic variants as proxy indicators (i.e. instrumental variables) for the exposure of interest (i.e. birth weight) and uses these genetic variants to assess the causal effect of the exposure on the outcome variable of interest (i.e. adult diseases) (Fig. 1) [29]. Because genetic variants are measured with high accuracy and capture long-term effect of the exposure, MR analysis results are often not susceptible to bias caused by measurement errors that are commonly encountered in randomised intervention studies [32]. In addition, because the two alleles of a genetic variant are randomly segregated during gamete formation and conception under the Mendel’s law and because such segregation is independent of many known or unknown confounders, MR analysis results are also less susceptible to reverse causation and confounding factors compared with other study designs [33]. As a result, MR has become a popular and cost-effective analysis tool for causal inference in observational studies, avoiding the need to record and control for all possible confounding factors present in the study.

**Fig. 1.**
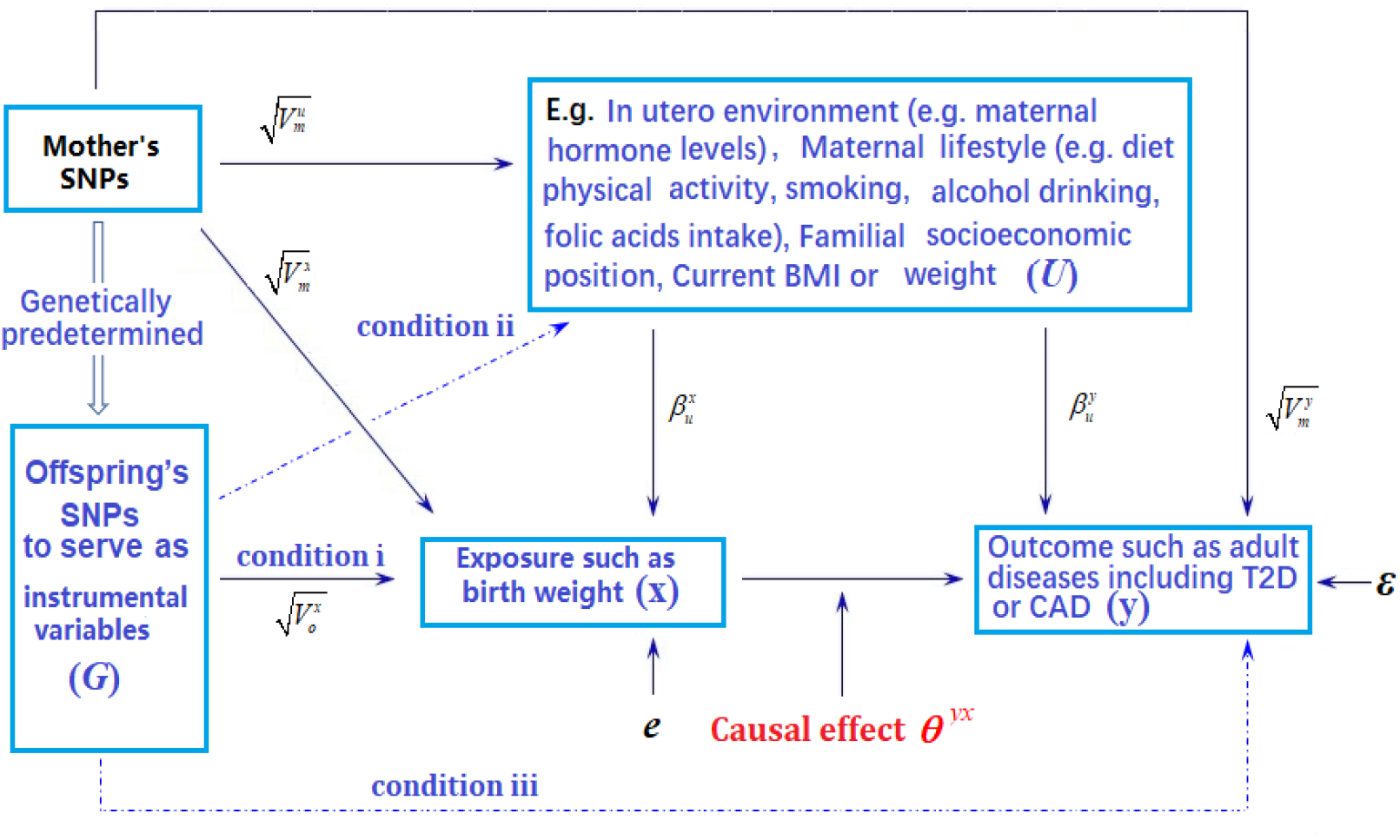
Graphical illustration of MR analysis. Arrows or dot lines represent the presence or absence of associations, respectively. The MR analysis estimates the causal effect of birth weight to adult disease risk in the presence of various measured and unmeasured confounding factors by carefully selecting SNPs that are associated with birth weight to serve as instrumental variables. Valid MR requires these selected SNPs to satisfy three conditions: selected SNPs are strongly associated with birth weight (condition **i**); selected SNPs are not associated with any known or unknown confounders that are associated with both birth weight and disease (condition **ii**); selected SNPs are independent of adult disease conditional on birth weight (condition **iii**). Note that the effects of instrumental variables (G) on the exposure of interest (x) may be indirect and mediated through mediator variables. Exemplary traits include BMI (body mass index), T2D (type 2 diabetes) and CAD (coronary artery disease). The notations in the figure are defined further in Text S4

Indeed, MR studies have been recently carried out to investigate the causal effect of birth weight on either CAD or T2D (e.g. [34, 35]), each with a relatively small sample size and subsequently a small set of valid instrumental variables. Unfortunately, for CAD, the causality result of birth weight does not hold in follow-up sensitivity analyses and is not robust with respect to the choice of statistical methods [34, 35]. For T2D, sensitivity analyses were not carried out in the study [36], and it was thus unclear, for example, whether instrumental variable outliers selected in the study may impact the estimation of the causal effect of birth weight. Here, we perform a large-scale MR study to comprehensively investigate the causal effects of birth weight on a total of 21 diseases and 38 complex traits in adulthood. Our results are validated with a wide range of sensitivity analyses and simulations to ensure result robustness.

## Materials and Methods

We present a brief overview of the analysis procedure with technical details provided in Text S1-Text S4.

### Data Sources

We first obtained summary statistics in terms of marginal effect size estimate of single nucleotide polymorphism (SNP) and its standard error on birth weight from the Early Growth Genetics (EGG) consortium study [37]. The EGG consortium study is the largest genome-wide association study (GWAS) to date on birth weight (a continuous trait) and contains association results for 16,245,523 genotyped and imputed SNPs based on up to 153,781 individuals collected from 35 studies (Table S1). Next, to examine the causal effect of birth weight on adult diseases, we collected summary statistics from corresponding GWASs for 21 diseases (Text S1). These diseases include advanced age-related macular degeneration (AMD) [38], Alzheimer’s disease [39], Parkinson’s disease [40], chronic kidney disease (CKD) [41], celiac disease [42], inflammatory bowel disease (IBD) [43], Crohn’s disease (CD) [43], ulcerative colitis (UC) [43], primary biliary cirrhosis (PBC) [44], primary sclerosing cholangitis (PSC) [45], systemic lupus erythematosus (SLE) [46], coronary artery disease (CAD) [47], myocardial infarction (MI) [47], type 2 diabetes (T2D) [36], rheumatoid arthritis (RA) [48], type 1 diabetes (T1D) [48], hypertension [48], ankylosing spondylitis (AS) [49], ischaemic stroke (IS) [49] and multiple sclerosis (MS) [49]. Finally, to identify complex traits that may mediate the causal effect of birth weight on any identified adult disease, we also obtained GWAS summary statistics for 38 complex traits in adulthood (Text S2). These traits include educational attainment (i.e. EduYears and College) [50], smoking behaviors [51], early growth traits [52], blood lipid traits [53], glycaemic and harmonic traits [54] and blood pressures [55]. With these data we performed MR analyses with series of sensitivity analyses. Detailed modeling assumptions and methodological considerations of MR analyses are described in Text S3.

### Selecting instruments for Mendelian randomisation analyses

We first selected 47 independent index genetic variants (Table 1) to serve as valid instrumental variables for birth weight based on the EGG consortium study [37] using the clumping procedure of the plink software (version v1.90b3.38) (Text S3) [56] following previous work [57]. Next, for each disease in turn, we relied on the corresponding GWAS and extracted summary statistics on the disease for the 47 index SNPs of birth weight. While our main MR analyses were performed using the above 47 SNPs as instrumental variables, to examine the robustness of the results, we also performed alternative MR analyses using a slightly different set of 48 SNPs. These 48 SNPs are presented in the original GWAS of birth weight [37] and that are independent, un-clumped index SNPs showing strong association with birth weight (*p* < 5.00E-8) (Table S2). For these index SNPs that do not have summary statistics in the corresponding disease, we either replaced them with proxy SNPs that are in high linkage disequilibrium (LD) with the index SNPs or imputed the summary statistics [58] for the index SNPs — both approaches yield similar results (Text S3). Afterwards, we excluded among them SNPs that show horizontal pleiotropic associations with the outcome to ensure the validity of MR analysis (Tables S3 and S4, which list effect estimates with or without excluding these potentially horizontal pleiotropic SNPs) [29, 59]. The number of instruments excluded varies for different diseases and ranges from 1 (e.g. for age-related macular degeneration) to 18 (e.g. for height) (Table S5). The final set of SNPs that are used as instruments differs across diseases and ranges from 23 (for Parkinson’s disease) to 47 (for Crohn’s disease) in our main MR analyses.

**Table 1.**
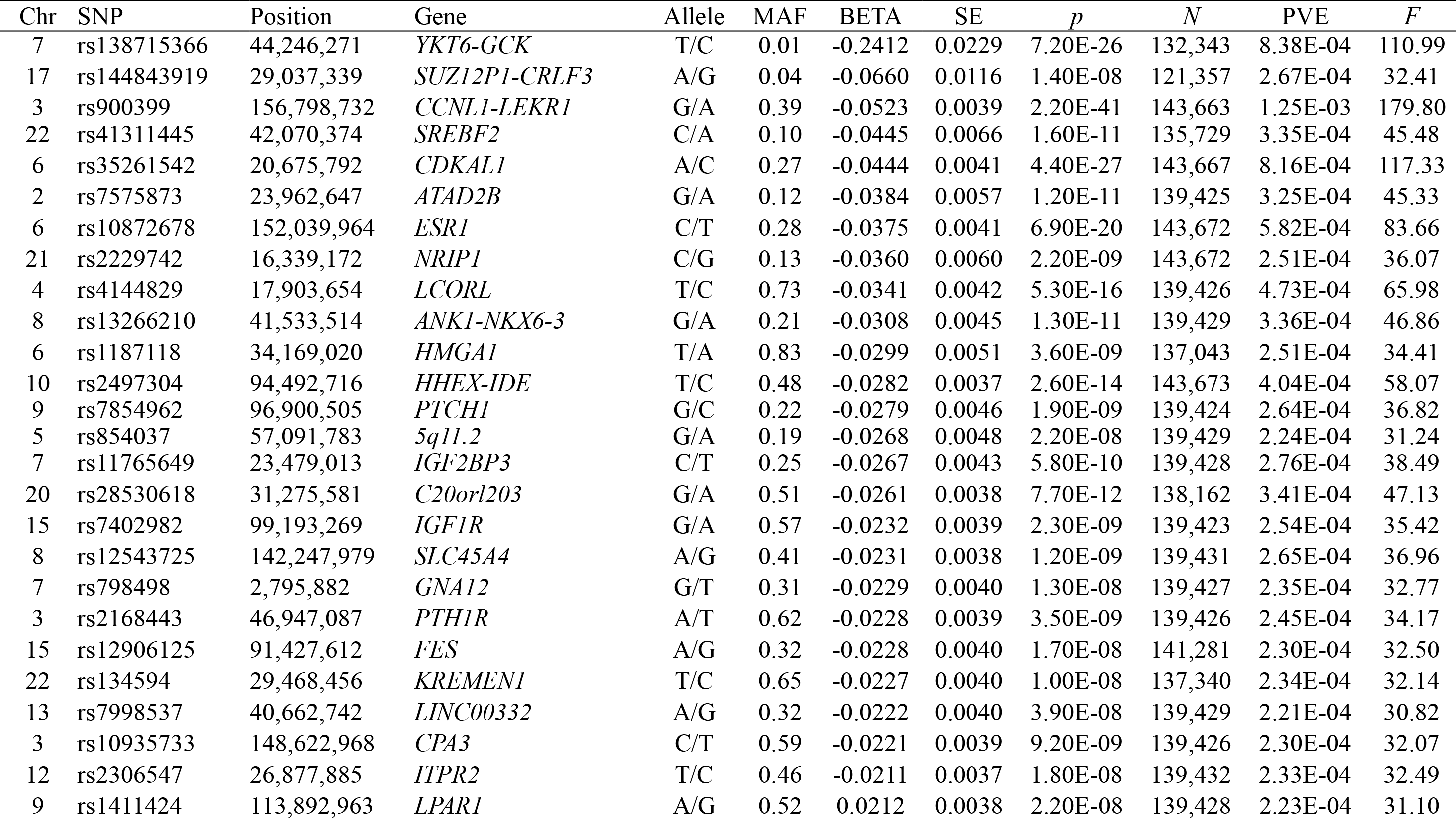
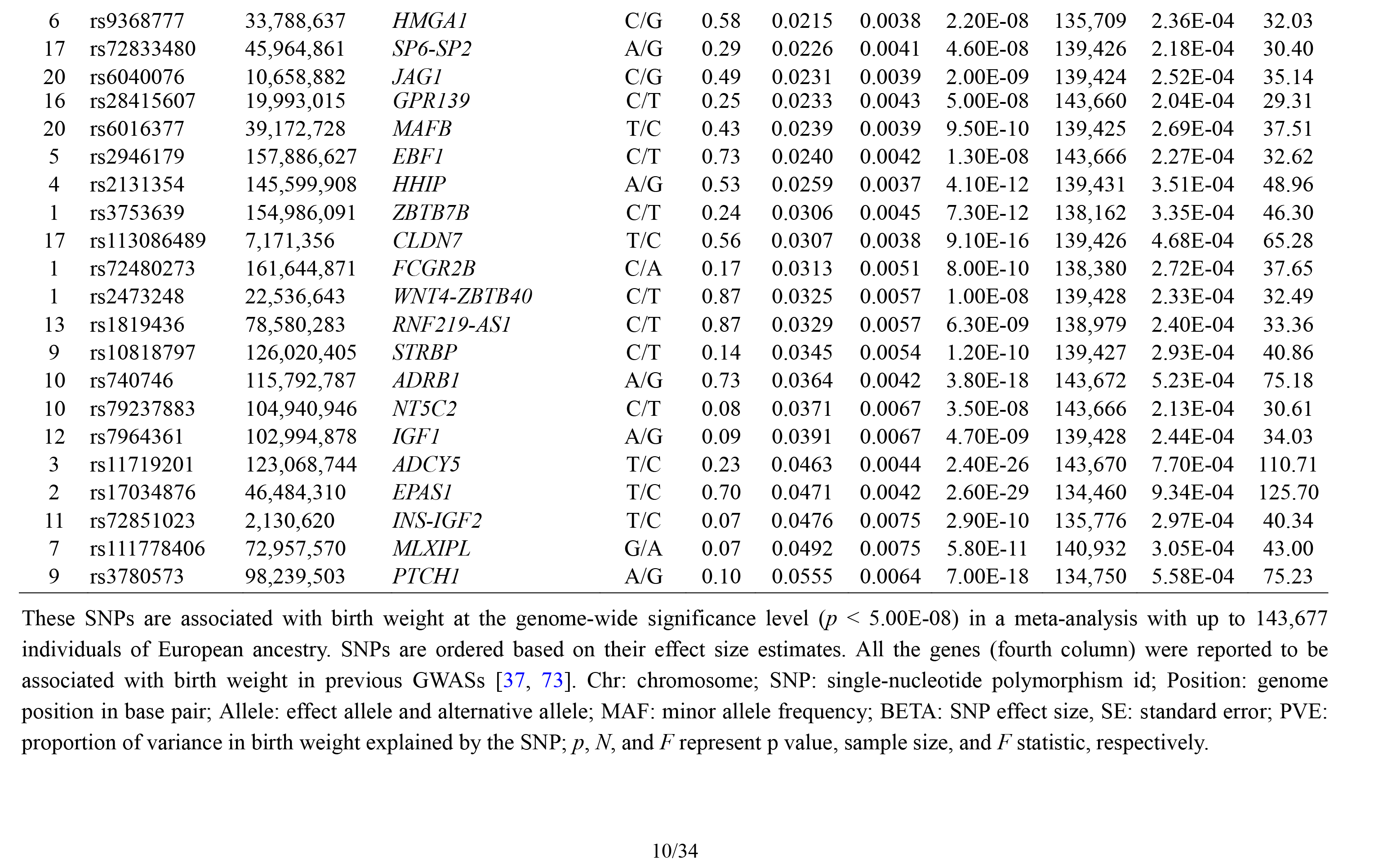
Summary information for the 47 autosomal SNPs that are used as instrumental variables in the MR analysis

| Chr | SNP | Position | Gene | Allele | MAF | BETA | SE | *p* | *N* | PVE | *F* |
| --- | --- | --- | --- | --- | --- | --- | --- | --- | --- | --- | --- |
| 7 | rs138715366 | 44,246,271 | *YKT6-GCK* | T/C | 0.01 | -0.2412 | 0.0229 | 7.20E-26 | 132,343 | 8.38E-04 | 110.99 |
| 17 | rs144843919 | 29,037,339 | *SUZ12P1-CRLF3* | A/G | 0.04 | -0.0660 | 0.0116 | 1.40E-08 | 121,357 | 2.67E-04 | 32.41 |
| 3 | rs900399 | 156,798,732 | *CCNL1-LEKRI* | G/A | 0.39 | -0.0523 | 0.0039 | 2.20E-41 | 143,663 | 1.25E-03 | 179.80 |
| 22 | rs41311445 | 42,070,374 | *SREBF2* | C/A | 0.10 | -0.0445 | 0.0066 | 1.60E-11 | 135,729 | 3.35E-04 | 45.48 |
| 6 | rs35261542 | 20,675,792 | *CDKAL1* | A/C | 0.27 | -0.0444 | 0.0041 | 4.40E-27 | 143,667 | 8.16E-04 | 117.33 |
| 2 | rs7575873 | 23,962,647 | *ATAD2B* | G/A | 0.12 | -0.0384 | 0.0057 | 1.20E-11 | 139,425 | 3.25E-04 | 45.33 |
| 6 | rs10872678 | 152,039,964 | *ESR1* | C/T | 0.28 | -0.0375 | 0.0041 | 6.90E-20 | 143,672 | 5.82E-04 | 83.66 |
| 21 | rs2229742 | 16,339,172 | *NRIP1* | C/G | 0.13 | -0.0360 | 0.0060 | 2.20E-09 | 143,672 | 2.51E-04 | 36.07 |
| 4 | rs4144829 | 17,903,654 | *LCORL* | T/C | 0.73 | -0.0341 | 0.0042 | 5.30E-16 | 139,426 | 4.73E-04 | 65.98 |
| 8 | rs13266210 | 41,533,514 | *ANK1-NKX6-3* | G/A | 0.21 | -0.0308 | 0.0045 | 1.30E-11 | 139,429 | 3.36E-04 | 46.86 |
| 6 | rs1187118 | 34,169,020 | *HMGA1* | T/A | 0.83 | -0.0299 | 0.0051 | 3.60E-09 | 137,043 | 2.51E-04 | 34.41 |
| 10 | rs2497304 | 94,492,716 | *HHEX-IDE* | T/C | 0.48 | -0.0282 | 0.0037 | 2.60E-14 | 143,673 | 4.04E-04 | 58.07 |
| 9 | rs7854962 | 96,900,505 | *PTCH1* | G/C | 0.22 | -0.0279 | 0.0046 | 1.90E-09 | 139,424 | 2.64E-04 | 36.82 |
| 5 | rs854037 | 57,091,783 | *5q11.2* | G/A | 0.19 | -0.0268 | 0.0048 | 2.20E-08 | 139,429 | 2.24E-04 | 31.24 |
| 7 | rs11765649 | 23,479,013 | *IGF2BP3* | C/T | 0.25 | -0.0267 | 0.0043 | 5.80E-10 | 139,428 | 2.76E-04 | 38.49 |
| 20 | rs28530618 | 31,275,581 | *C20orf203* | G/A | 0.51 | -0.0261 | 0.0038 | 7.70E-12 | 138,162 | 3.41E-04 | 47.13 |
| 15 | rs7402982 | 99,193,269 | *IGF1R* | G/A | 0.57 | -0.0232 | 0.0039 | 2.30E-09 | 139,423 | 2.54E-04 | 35.42 |
| 8 | rs12543725 | 142,247,979 | *SLC45A4* | A/G | 0.41 | -0.0231 | 0.0038 | 1.20E-09 | 139,431 | 2.65E-04 | 36.96 |
| 7 | rs798498 | 2,795,882 | *GNAI2* | G/T | 0.31 | -0.0229 | 0.0040 | 1.30E-08 | 139,427 | 2.35E-04 | 32.77 |
| 3 | rs2168443 | 46,947,087 | *PTH1R* | A/T | 0.62 | -0.0228 | 0.0039 | 3.50E-09 | 139,426 | 2.45E-04 | 34.17 |
| 15 | rs12906125 | 91,427,612 | *FES* | A/G | 0.32 | -0.0228 | 0.0040 | 1.70E-08 | 141,281 | 2.30E-04 | 32.50 |
| 22 | rs134594 | 29,468,456 | *KREMEN1* | T/C | 0.65 | -0.0227 | 0.0040 | 1.00E-08 | 137,340 | 2.34E-04 | 32.14 |
| 13 | rs7998537 | 40,662,742 | *LINC00332* | A/G | 0.32 | -0.0222 | 0.0040 | 3.90E-08 | 139,429 | 2.21E-04 | 30.82 |
| 3 | rs10935733 | 148,622,968 | *CPA3* | C/T | 0.59 | -0.0221 | 0.0039 | 9.20E-09 | 139,426 | 2.30E-04 | 32.07 |
| 12 | rs2306547 | 26,877,885 | *ITPR2* | T/C | 0.46 | -0.0211 | 0.0037 | 1.80E-08 | 139,432 | 2.33E-04 | 32.49 |
| 9 | rs1411424 | 113,892,963 | *LPAR1* | A/G | 0.52 | 0.0212 | 0.0038 | 2.20E-08 | 139,428 | 2.23E-04 | 31.10 |
| 6 | rs9368777 | 33,788,637 | *HMGA1* | C/G | 0.58 | 0.0215 | 0.0038 | 2.20E-08 | 135,709 | 2.36E-04 | 32.03 |
| 17 | rs72833480 | 45,964,861 | *SP6-SP2* | A/G | 0.29 | 0.0226 | 0.0041 | 4.60E-08 | 139,426 | 2.18E-04 | 30.40 |
| 20 | rs6040076 | 10,658,882 | *JAG1* | C/G | 0.49 | 0.0231 | 0.0039 | 2.00E-09 | 139,424 | 2.52E-04 | 35.14 |
| 16 | rs28415607 | 19,993,015 | *GPR139* | C/T | 0.25 | 0.0233 | 0.0043 | 5.00E-08 | 143,660 | 2.04E-04 | 29.31 |
| 20 | rs6016377 | 39,172,728 | *MAFB* | T/C | 0.43 | 0.0239 | 0.0039 | 9.50E-10 | 139,425 | 2.69E-04 | 37.51 |
| 5 | rs2946179 | 157,886,627 | *EBF1* | C/T | 0.73 | 0.0240 | 0.0042 | 1.30E-08 | 143,666 | 2.27E-04 | 32.62 |
| 4 | rs2131354 | 145,599,908 | *HHIP* | A/G | 0.53 | 0.0259 | 0.0037 | 4.10E-12 | 139,431 | 3.51E-04 | 48.96 |
| 1 | rs3753639 | 154,986,091 | *ZBTB7B* | C/T | 0.24 | 0.0306 | 0.0045 | 7.30E-12 | 138,162 | 3.35E-04 | 46.30 |
| 17 | rs113086489 | 7,171,356 | *CLDN7* | T/C | 0.56 | 0.0307 | 0.0038 | 9.10E-16 | 139,426 | 4.68E-04 | 65.28 |
| 1 | rs72480273 | 161,644,871 | *FCGR2B* | C/A | 0.17 | 0.0313 | 0.0051 | 8.00E-10 | 138,380 | 2.72E-04 | 37.65 |
| 1 | rs2473248 | 22,536,643 | *WNT4-ZBTB40* | C/T | 0.87 | 0.0325 | 0.0057 | 1.00E-08 | 139,428 | 2.33E-04 | 32.49 |
| 13 | rs1819436 | 78,580,283 | *RNF219-AS1* | C/T | 0.87 | 0.0329 | 0.0057 | 6.30E-09 | 138,979 | 2.40E-04 | 33.36 |
| 9 | rs10818797 | 126,020,405 | *STRBP* | C/T | 0.14 | 0.0345 | 0.0054 | 1.20E-10 | 139,427 | 2.93E-04 | 40.86 |
| 10 | rs740746 | 115,792,787 | *ADRB1* | A/G | 0.73 | 0.0364 | 0.0042 | 3.80E-18 | 143,672 | 5.23E-04 | 75.18 |
| 10 | rs79237883 | 104,940,946 | *NT5C2* | C/T | 0.08 | 0.0371 | 0.0067 | 3.50E-08 | 143,666 | 2.13E-04 | 30.61 |
| 12 | rs7964361 | 102,994,878 | *IGF1* | A/G | 0.09 | 0.0391 | 0.0067 | 4.70E-09 | 139,428 | 2.44E-04 | 34.03 |
| 3 | rs11719201 | 123,068,744 | *ADCY5* | T/C | 0.23 | 0.0463 | 0.0044 | 2.40E-26 | 143,670 | 7.70E-04 | 110.71 |
| 2 | rs17034876 | 46,484,310 | *EPAS1* | T/C | 0.70 | 0.0471 | 0.0042 | 2.60E-29 | 134,460 | 9.34E-04 | 125.70 |
| 11 | rs72851023 | 2,130,620 | *INS-IGF2* | T/C | 0.07 | 0.0476 | 0.0075 | 2.90E-10 | 135,776 | 2.97E-04 | 40.34 |
| 7 | rs111778406 | 72,957,570 | *MLXIPL* | G/A | 0.07 | 0.0492 | 0.0075 | 5.80E-11 | 140,932 | 3.05E-04 | 43.00 |
| 9 | rs3780573 | 98,239,503 | *PTCH1* | A/G | 0.10 | 0.0555 | 0.0064 | 7.00E-18 | 134,750 | 5.58E-04 | 75.23 |

These SNPs are associated with birth weight at the genome-wide significance level ( $p < 5.00E-08$ ) in a meta-analysis with up to 143,677 individuals of European ancestry. SNPs are ordered based on their effect size estimates. All the genes (fourth column) were reported to be associated with birth weight in previous GWASs [37, 73]. Chr: chromosome; SNP: single-nucleotide polymorphism id; Position: genome position in base pair; Allele: effect allele and alternative allele; MAF: minor allele frequency; BETA: SNP effect size, SE: standard error; PVE: proportion of variance in birth weight explained by the SNP; $p$ , $N$ , and $F$ represent $p$ value, sample size, and $F$ statistic, respectively.

### Mendelian randomisation, sensitivity analyses and multivariable analyses

The main statistical details for data analyses are provided in Text S3. For either set of the instrumental variables, for each SNP in turn we estimated the proportion of phenotypic variance explained (PVE) by the SNP using summary statistics and further computed *F* statistic to ensure strong instruments. We then performed MR analyses using both the fixed-effects version and the random-effects version of the inverse-variance weighted (IVW) methods [60-62] to estimate and test the causal effects of birth weight on each of the 21 adult diseases. Compared with the fixed-effects version, the random-effects version of IVW accounts for causal effect size heterogeneity across instruments and often yield more conservative results. Besides applying the random-effects version of IVW, we also employ both the Q and I^2^ statistics to directly measure effect size heterogeneity in the data [63].

For each disease identified to be causally affected by birth weight in the MR analyses, we performed a series of sensitivity analyses to ensure results robustness. Specifically, we performed a median-based MR analysis to guard against outlying instruments [64]. We conducted a leave-one-out (LOO) cross validation analysis to directly examine potential instrument outliers [57]. We carried out MR-Egger regression to examine the assumption of directional pleiotropic effects [65, 66]. To examine the potential influence of maternal genetic effects, we performed additionally sensitivity analyses by excluding SNPs that affect birth weight through maternal effects [21, 67, 68]. In addition, we carried out simulations to examine the potential impact of maternal effects on causal effect estimations [21] (Text S4). We performed reverse causal inference to examine the possible reverse causality from diseases to birth weight. We also applied a recently developed analysis method iMAP [69] to jointly model all genome-wide SNPs to provide supportive evidence on the directionality of the causal relationship between birth weight and these identified diseases.

Finally, we investigated whether any of the 38 complex traits may mediate the causal effect of birth weight on the identified adult diseases. To do so, we first performed MR analysis to examine whether birth weight causally affect any of the 38 complex traits. In particular, for each of the 38 traits in turn, we extracted summary statistics from the corresponding GWAS for the 47 instrumental variables of birth weight. We replaced missing SNPs with proxy ones when necessary and applied the IVW methods following the same procedure as described above. Next, we further performed a multivariable MR analysis [70-72] for each pair of identified trait and disease to investigate whether any of these complex traits may mediate the causal effect of birth weight on the identified disease. The multivariable MR analysis allows us to estimate and test both the direct effect of birth weight on the disease and the indirect effect of birth weight on the disease through the complex trait [72].

## Results

### Mendelian randomisation identifies three adult diseases that are causally affected by birth weight

We first selected a set of 47 SNPs from a large-scale GWAS for birth weight based on 143,677 individuals to serve as instrumental variables for birth weight (Table 1 and Fig. S1). We examined the strength of these instrumental variables using *F* statistic based on the EGG GWAS discovery sample of birth weight following [57] (details in Text S3). For the 47 instrumental variables, their *F* statistics individually range from 29.36 to 179.83 (Table 1) with an overall *F* statistic of 49.22 for all 47 instruments. These values are all above the usual threshold of 10, suggesting that the selected genetic variants have sufficiently strong effect sizes to be used as instrumental variables and that weak instrument bias is unlikely to occur in our analysis.

Across 21 diseases, with either the random-effects (Fig. 2A) or the fixed-effects (Fig. 2B) IVW approaches (see also Table S3). Note that we displayed the causal effects of *lower* birth weight instead of birth weight in all figures and tables throughout the text by supplying a negative sign on the estimated birth weight effect. We found that lower birth weight is causally associated with increased risks for three diseases after Bonferroni correction (i.e. *p*-value threshold of 0.05/21 = 2.38E-3). These three diseases include coronary artery disease (CAD), myocardial infarction (MI) [47], and type 2 diabetes (T2D; both in terms of the original T2D status and in terms of T2D_BMI which represents the T2D status after adjusting for BMI) [36]. Both the random-effects and fixed effects IVW methods yield identical estimates for the causal effect sizes, but the former generates slightly wider confidence intervals than the later as one would expect. Because the random-effects IVW properly accounts for causal effect heterogeneity estimated using each of the 47 instruments and because we indeed identified such heterogeneity (Table S3; e.g. p values based on Q statistic are 7.41E-1, 1.42E-2, 1.40E-4 and 1.92E-2, and the I^2^ statistics are 0%, 33.1%, 48.9% and 31.8% for CAD, MI, T2D and T2D_BMI, respectively), we choose to mainly present our results from the random-effects IVW analysis in the following main text.

**Fig. 2.**
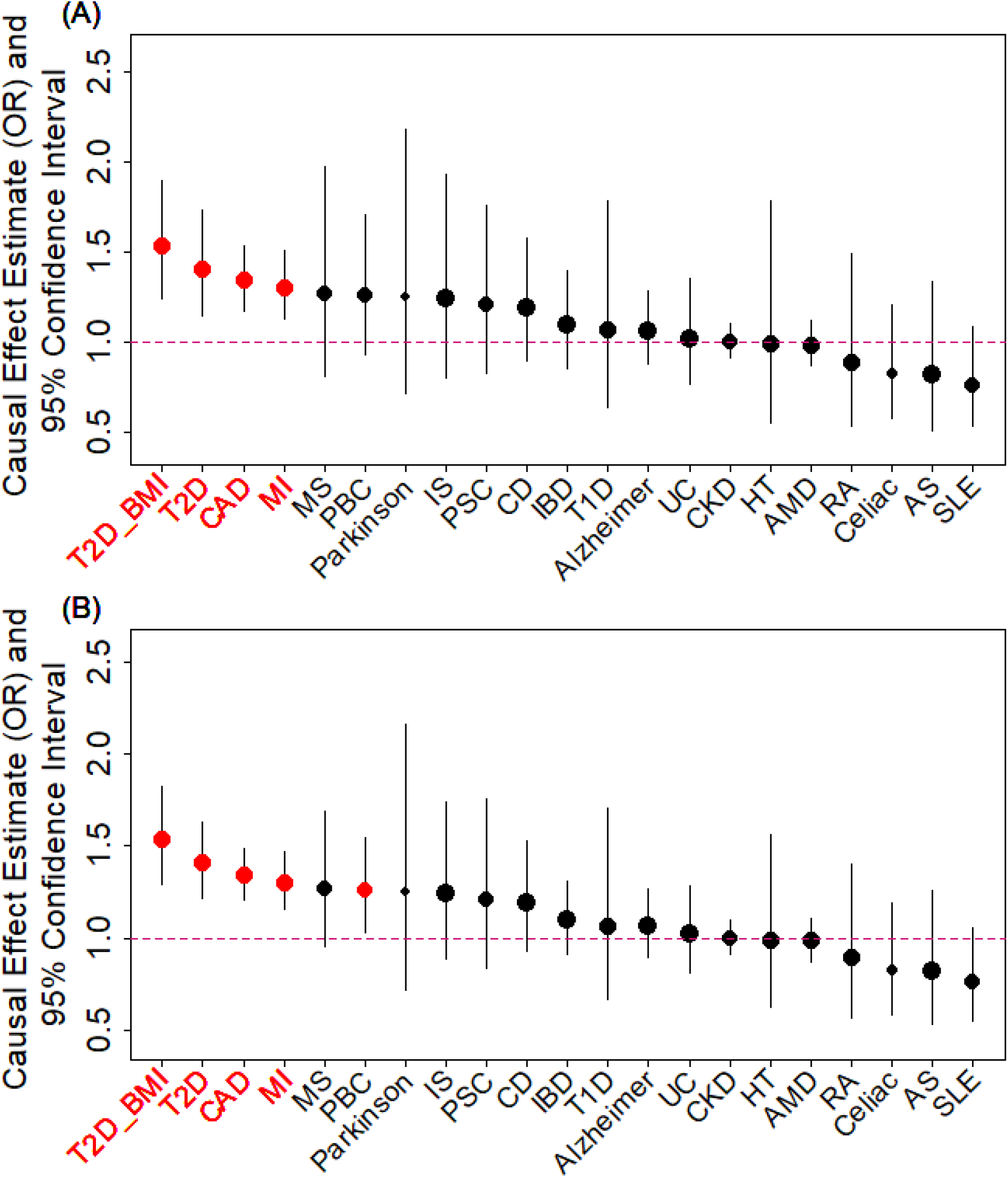
Causal effect estimates and 95% confidence intervals for lower birth weight on 21 diseases based on either the random-effects inverse variance weighted (IVW) method (**A**) or the fixed-effects IVW method (**B**). Diseases are ordered based on their causal effect estimates. Estimations are carried out using both index SNPs and proxy SNPs. In both panels, dot size is proportional to the number of instrumental variables used for the given disease while dot color represents significance (*p* < 0.05 are highlighted in red). Disease names (x-axis) are further highlighted in red if the causal effects are significant after Bonferroni correction (*p* < 0.05/21)

For each of the three diseases, we compute the odds ratio (OR) of the disease for one unit decrease of birth weight, where the unit is defined as the standard deviation of birth weight, estimated to be 488 grams across 35 studies in the original meta-analysis (Table S1) [37]. We found that a unit lower birth weight is causally associated with an increased CAD risk with an estimated OR of 1.34 [95% confidence interval (CI) 1.17 - 1.53, *p* = 1.54E-5]; a unit lower birth weight is causally associated with an increased MI risk with an estimated OR of 1.30 (95% CI 1.13 - 1.51, *p* = 3.31E-4); a unit lower birth weight is also causally associated with an increased T2D risk, with an estimated OR of 1.41 (95% CI 1.15 - 1.73, *p* = 1.11E-3) for the original T2D, and with an estimated OR of 1.54 (95% CI 1.25 - 1.89, *p* = 6.07E-5) for the BMI-adjusted T2D (i.e. T2D_BMI).

Finally, consistent with the fetal origins of adult diseases hypothesis, the causal effects of lower birth weight on most of the diseases investigated are estimated to be positive [14 out of 21 (66.7%), Table S3], though most of these estimates are not statistically significantly different from one. In addition, the estimated causal effects of lower birth weight on five diseases (ischaemic stroke, multiple sclerosis, Parkinson’s disease, primary biliary cirrhosis, and primary sclerosing cholangitis) in addition to the three diseases mentioned in the above paragraph are above OR of 1.2, though these estimates came with large standard errors. Power simulation results based on parameters estimated in the MR analysis also suggest that the nonsignificant results for the remaining diseases may be due to a lack of statistical power (Table S3). The lack of power for the remain diseases suggest that a lack of association between birth weight and these diseases should not be over-interpreted and that larger sample sizes are needed to elucidate the causal effects of birth weight on these diseases.

### Mendelian randomisation results are robust with respect to instrument outliers and the choice of instrumental variables

We examine the causal relationship between birth weight and the three diseases (CAD, MI, T2D, and T2D_BMI) in details here. We first display the causal effects of lower birth weight for each of the three diseases estimated using individual instrumental variables in Fig. 3. We also plot the SNP effect sizes on birth weight versus the effect sizes on these diseases in Fig. 4. One SNP, rs138715366, appears to be an outlier for all these traits. rs138715366 has a low minor allele frequency (MAF = 0.89%), is located within the intronic regions of the gene *YKT6-GCK* on Chr 7 and has the largest effect size on birth weight among all instrument variables (= −0.24; with 95% CI −0.20 - −0.29; *p* = 7.20E-26; Fig. S1 and Table 1). In addition, another SNP, rs144843919, also appears to be a potential outlier for CAD. The effect size of rs144843919 on birth weight is estimated to −0.066 (95% CI −0.09 - −0.04, *p* = 1.40E-8). However, as we will show in the next paragraph, neither SNP has substantial influence on the estimation of the causal effects.

**Fig. 3.**
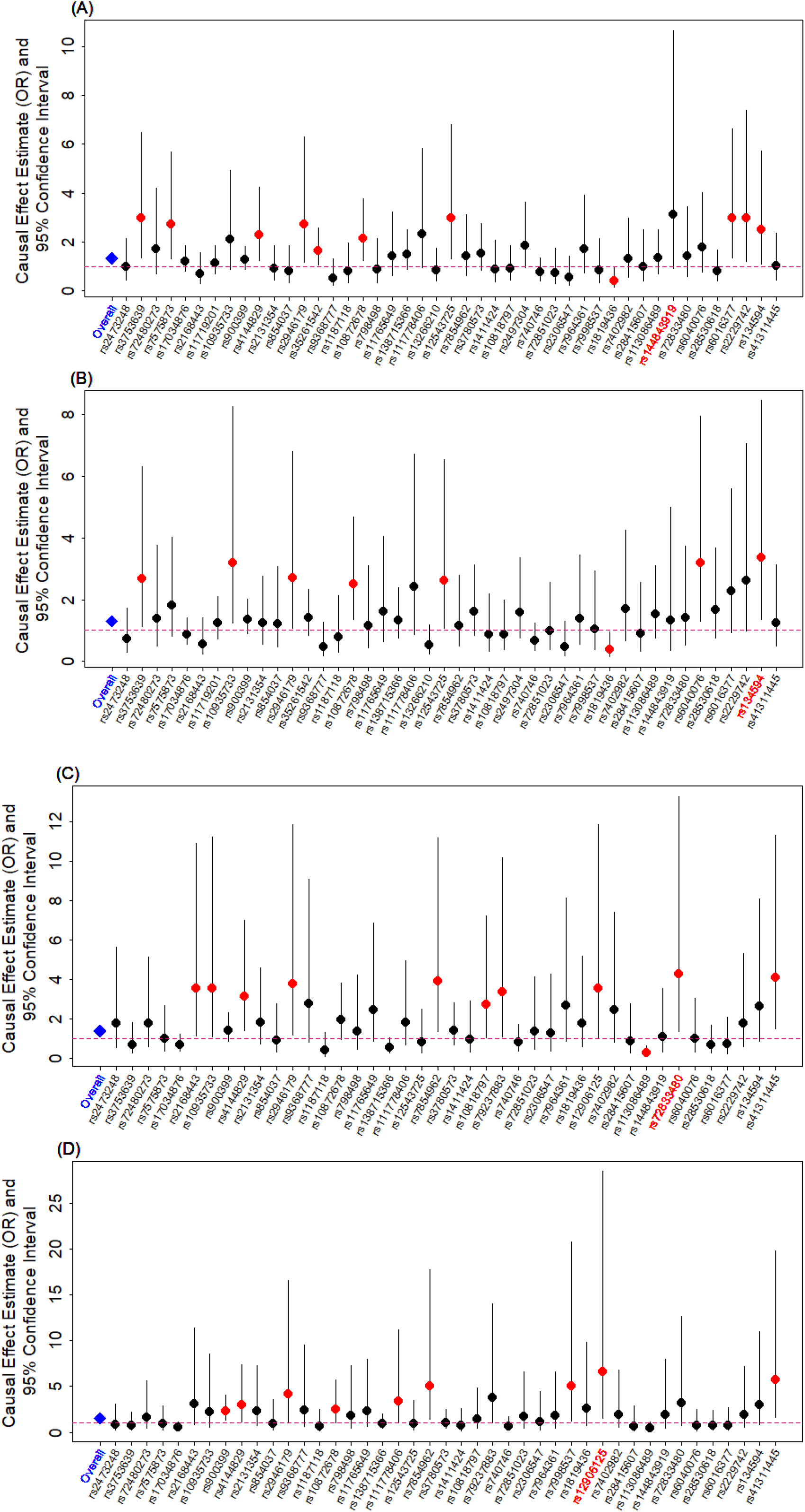
Causal effect estimates and 95% confidence intervals for lower birth weight on (**A**) CAD, (**B**) MI, (**C**) T2D, and (**D**) T2D_BMI. Estimations are carried out either using all SNPs (first column on x-axis) or using individual SNPs (the remaining columns on x-axis) based on Equation (14) in Text S3. Dot size is proportional to the effect size estimates while dot color represents significance (*p* < 0.05 are highlighted in red). SNP that yields the largest causal effect estimate is also highlighted in red (x-axis)

**Fig. 4.**
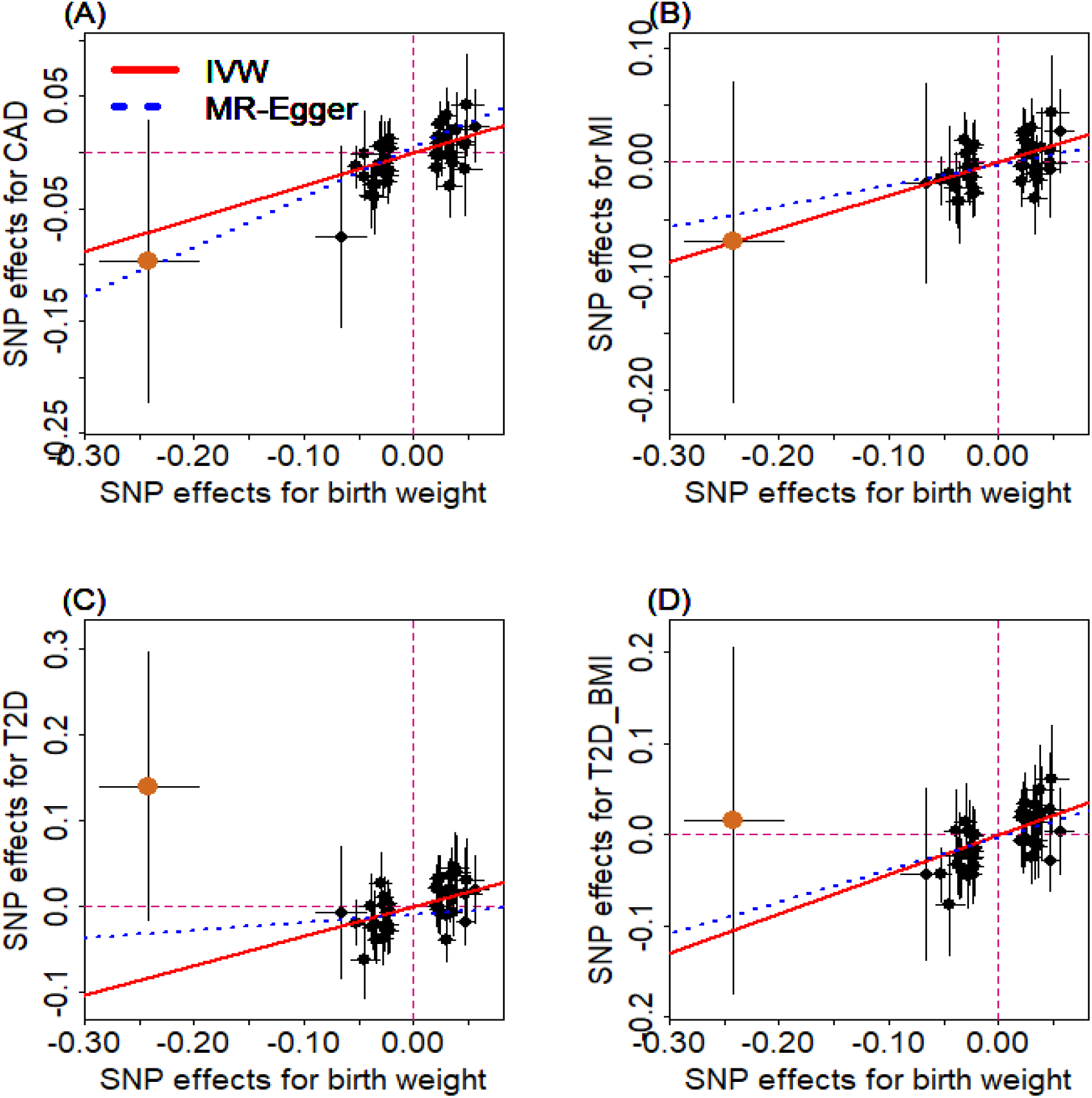
Relationship between the effect size estimates on lower birth weight (x-axis) and the effect size estimates on diseases (y-axis) for the 47 SNPs that serve as instrumental variables. Examined diseases include (**A**) CAD, (**B**) MI, (**C**) T2D, and (**D**) T2D_BMI. 95% confidence intervals for the estimated SNP effect sizes on disease are shown as vertical black lines, while the 95% confidence intervals for the estimated SNP effect sizes on birth weight are shown as horizontal black lines. The vertical and horizontal red dotted lines represent zero effects. The slope of fitted lines represents the estimated the casual effects of birth weight on the corresponding disease obtained using either the IVW methods (red solid lines) or the MR-Egger regression (blue dotted lines). SNP outlier rs13875366 (chocolate dot) was not included in MR-Egger regression to avoid outlier influence. Due to the inclusion of an intercept in the MR-Egger regression, the fitted lines by MR-Egger regression (blue dotted lines) do not necessarily pass the origin

The LOO analysis results are stable and demonstrate that no single instrumental variable substantially influences the estimation of the casual effects of birth weight on CAD, MI, T2D, or T2D_BMI (Fig. S2). For example, after removing rs138715366, the ORs for a unit decrease of birth weight are estimated to be 1.34 (95% CI 1.20 - 1.49, *p* = 9.70E-8) for CAD, 1.30 (95% CI 1.15 - 1.47, *p* = 1.66E-5) for MI, 1.48 (95% CI 1.27 - 1.71*,p* = 2.66E-7) for T2D and 1.57 (95% CI 1.32 - 1.87*,p* = 3.95E-7) for T2D_BMI, almost identical to the ORs estimated using all these instrumental variables together (Fig. 2).

Our primary results described in the previous section are based on using 47 instrumental variables. For certain diseases, summary statistics for some of the 47 index SNPs are unavailable. In these cases, we have used proxy SNPs that are in high LD using a certain correlation threshold. We found that our results are robust with respect to various correlation thresholds to obtain these proxy SNPs (Fig. S3). Besides using proxy SNPs, we imputed summary statistics for the unavailable index SNPs and performed analysis using all index SNPs. Results with imputed summary statistics remain similar (Fig. S4). We also performed analysis using only part of the 47 index SNPs that are available for the given disease, without using any proxy SNPs or imputation; we again obtained consistent results (Fig. S5). Finally, besides the analysis using clumped SNPs, we performed an alternative analysis by using 48 un-clumped instrumental variables (Table S2) that are presented in the original meta-analysis study [37]. Again, the alternative analysis results are largely similar to those in our main analyses (Fig. S6 and Fig. S7).

### Various sensitivity analyses further validate the main Mendelian randomisation results

We performed sensitivity analyses to complement our main MR analysis results obtained by using the IVW approaches. First, to guard against the possibility that some instruments are invalid, we conducted a MR analysis using the weighted median method [64] for CAD, MI, T2D and T2D_BMI. The weighted median estimate approach yields qualitatively similar results as our main analysis (Text S3), suggesting that invalid instruments unlikely bias our main results.

To guard against the possibility that the used instruments may display horizontal pleiotropy and thus bias causal effect estimation, we performed the MR-Egger regression [65, 66] for the four traits. The results from the MR-Egger regression analysis are again consistent with our main results (Text S3, see also Fig. 2). In addition, none of the intercepts from MR-Egger regression are significantly deviated from zero: they are estimated to be 0.005 (95% CI −0.010 - 0.020, *p* = 0.515) for CAD, −0.003 (95% CI −0.019 - 0.013, *p* = 0.756) for MI, −0.010 (95% CI −0.031 - 0.012, *p* = 0.383) for T2D and −0.003 (95% CI −0.026 - 0.019, *p* = 0.783) for T2D_BMI, respectively. Moreover, funnel plots also display symmetric pattern of effect size variation around the point estimate (Fig. S8). Together, MR-Egger regression results and funnel plots suggest that horizontal pleiotropy unlikely bias our results.

One of the main difficulties in causal inference is to distinguish causality from reverse causality [60]. Because of the time order and the fact that birth weight precedes adult diseases, the issue of reverse causation is unlikely a concern in our study. Nevertheless, to guard against the small possibility that our results are driven by reverse causality, we performed IVW analysis in the reverse direction to examine the causal effects of CAD, MI, T2D, or T2D_BMI on birth weight. Results show that there are no reverse causal associations between any of the four traits and birth weight as one would expect (Fig. S10).

To complement the MR analysis, we also performed analysis using the recently developed iMAP method [69]. iMAP analyzes a pair of traits jointly and borrows information across all genome-wide SNPs to provide additional evidence regarding to the causal relationship between the two traits. In particular, iMAP estimates the proportion of SNPs associated with one trait that is also associated with the other. By estimating such proportions, iMAP has the potential to provide evidence supporting potentially directional causality between the two analyzed traits [69, 74]. Here, we applied iMAP to analyze birth weight and each of the four traits at a time. We estimated the proportions of SNPs associated with birth weight that are also associated with CAD, MI, T2D, and T2D_BMI to be 0.125, 0.134, 0.452, and 0.472, respectively. In contrast, the proportion of SNPs associated with CAD, MI, T2D, and T2D_BMI that is also associated with birth weight are only 0.053, 0.029, 0.211, and 0.130, respectively. The asymmetrical probabilities estimated from iMAP suggest that SNPs associated with the birth weight are also more likely associated with the disease than the other way around. Therefore, iMAP provides additional genome-wide evidence supporting the causal effects of birth weight on the identified diseases.

### Causal effects of birth weight on the three identified diseases are not mediated through other complex traits

We explored the causal pathways through which birth weight causally affect the adult diseases. To do so, we obtained 35 quantitative traits and 3 binary traits that may mediate the causal effects of birth weight onto diseases (Text S2). For each trait in turn, we estimated the casual effect of birth weight on the trait using all available instruments using IVW (Fig. S11 and Table S4). Among all examined traits, we only identified adult weight to be causally affected by birth weight based on the Bonferroni adjusted significance threshold (*p* < 0.05/38) in both the fixed-effects and random-effects IVW analyses. In particular, the causal effect of birth weight on adult weight is estimated to be 0.36 in the fixed-effects IVW analysis (95% CI 0.17 - 0.55, *p* = 1.77E-4) and is estimated to be 0.36 in the fixed-effects IVW analysis (95% CI 0.12 - 0.60, *p* = 3.18E-3).

The lack of significant causal effects of birth weight on most examined complex traits are consistent with the lack of significant causal effects of birth weight on some of the examined diseases described in the earlier section. For example, we found that birth weight is not causally associated with both systolic blood pressure (SBP) and diastolic blood pressure (DBP) (estimated causal effect on SBP is 0.35, 95% CI −0.10 - 0.79, *p* = 0.127; estimated causal effect on DBP is 0.27, 95% CI −0.17 - 0.71, *p* = 0. 233) by random-effects IVW analysis. The lack of causal association between birth weight and blood pressure is consistent with our earlier result on a lack of detectable causal association between birth weight and hypertension. In addition, the lack of causal association between birth weight and many complex traits suggests that the causal effects of birth weight on CAD, MI or T2D are unlikely to be mediated by blood pressures or many other complex traits, which is further confirmed by the following multivariable regression.

To examine the possibility that some complex traits (e.g. adult weight, BMI, blood pressures, blood lipids, or hypertension) may mediate the causal effect of birth weight on each of the four traits (i.e. CAD, MI, T2D and T2D_BMI), we performed a comprehensive multivariable MR analysis for all the 38 complex traits (Text S2) [72]. The results provide the evidence that those complex traits are not likely to be the mediators of birth weight and support the conclusion that birth weight is an independent casual risk factor for CAD, MI, and T2D (Table S6). For example, the estimated direct effect for a unit lower birth weight for CAD, MI, T2D and T2D_BMI are 1.28 (95% CI 1.09 - 1.50, *p* = 4.57E-3), 1.28 (95% CI 1.07 - 1.52, *p* = 8.48E-3), 1.66 (95% CI 1.17 - 2.37, *p* = 6.84E-3) and 1.73 (95% CI 1.19 - 2.51, *p* = 6.28E-3). The estimated indirect effect of birth weight on CAD, MI, T2D and T2D_BMI are 1. 05 (95% CI 0.85 - 1.30, *p* = 0.653), 1.02 (95% CI 0.84 - 1.23, *p* = 0.852), 0.85 (95% CI 0.56 - 1.28, *p* = 0.432) and 0.89 (95% CI 0.58 - 1.37, *p* = 0.606), respectively. Therefore, the lack of detectable indirect effect suggests that adult weight unlikely mediate the causal effect of birth weight on any of the three diseases.

### Special sensitivity analyses to examine the influence of maternal effects on the main Mendelian randomisation results

Finally, we performed two additional sensitivity analyses to examine the influence of maternal effects on MR results (details available in the Text S4). First, we excluded among the set of 47 instruments those instruments that may potentially exhibit maternal effects on birth weights relying on a recent GWAS of maternal effects on birth weights [67]. We deleted a total of ten instruments, and with the remaining 37 instruments, we estimated the ORs (again, after removing the potentially pleiotropic instruments as done above) for a unit decrease in birth weight to be 1.37 (95% CI 1.17 - 1.61, *p* = 8.18E-5) for CAD, 1.31 (95% CI 1.10 - 1.55, *p* = 1.90E-3) for MI, 1.42 (95% CI 1.11 - 1.80, *p* = 4.62E-3) for T2D, and 1.41 (95% CI 1.10 - 1.80, *p* = 6.35E-3) for T2D_BMI, respectively. The results are consistent with the main results, suggesting that maternal effects unlikely bias our estimates. Second, we performed simulations to evaluate the extent to which the maternal effects may influence the birth-weight causal effect estimation in MR (Fig. S12) [21]. The simulation results (Fig. S13) show that the causal effects of birth weight are approximately unbiased when the maternal effect is in a reasonable range [37, 67] (e.g. each instrument has a maternal effect that explains 0.1% or 0.01% of phenotypic variance). Only when the maternal effect is unrealistically strong (e.g. each explains 1% or 10% of phenotypic variance), then, as one would expect, the causal effect estimates can be slightly biased upward. The approximate unbiasedness results in simulations also suggest that our main MR results are unlikely biased by realistic maternal effects.

## Discussion

### A summary of our Mendelian randomisation analyses

We have investigated the fetal origins of adult diseases hypothesis by performing a series of comprehensive MR analyses to examine the causal effects of birth weight on 21 adult diseases and 38 other complex traits. Our study relies on summary statistics obtained from 37 GWASs with sample sizes ranging from 4,798 (for rheumatoid arthritis [48]) to 339,224 (for BMI [75]), thus representing one of the largest and most comprehensive MR analyses performed on birth weight to date. The large sample size used in our study allows us to fully establish an inverse causal relationship between birth weight and three adult diseases that include CAD, MI and T2D. These inferred causal relationships are robust with respect to the selection of instrumental variables and to the choice of statistical methods, and are carefully validated in the present study through various sensitive analyses. In addition, our analysis also suggests that the lack of causality evidence between birth weight and the other diseases may be partly due to a lack of statistical power resulting from relatively small sample sizes for the remaining diseases. Finally, we investigate the possibility that any of the analyzed 38 complex traits may mediate the causal effects of birth weight on CAD, MI or T2D. Overall, our study provides important causality evidence supporting the fetal origins hypothesis for three adult diseases and suggests that increasing sample size is likely needed to reveal causal effects of birth weight for the other disorders.

### Comparison of our findings with those in previous studies

Our causality results are consistent with some of the previous association results obtained using standard logistic regressions. For example, we have estimated the OR of T2D per 488g lower of birth weight to be 1.41, which is very close to a previous meta-analysis estimate obtained using logistic regression where the OR of T2D per 500g lower of birth weight is estimated to be 1.47 [13]. We have estimated the OR of CAD per 488g lower of birth weight to be 1.34, which is also close to that obtained from a birth cohort study where the OR of CAD for a 500g decrease in birth weight is estimated to be 1.27 [10]. Our conclusions of T2D and CAD here are also consistent with those previously derived by a genetic risk score regression [34] and a similar MR analysis [68]. In addition, our results suggest that the inverse causal associations of birth weight with CAD, MI, or T2D are not likely mediated by other risk factors such as blood pressures or adult weight, again in line with previous studies [19]. Nevertheless, we also acknowledge that our results may appear to be inconsistent with those in [68] in terms of detecting the causal effects of birth weight on LDL, BMI and 2-hours glucose. However, for LDL and BMI, we note that our results are based on a more stringent p-value significance threshold adjusted by Bonferroni correction (to adjust for the multiple traits examined). Our results of birth weight on LDL or BMI are indeed marginally significant based on the normal p-value threshold of 0.05 with expected effect direction (i.e. negative effect on LDL and positive effect on BMI), and are thus consistent with [68]. For 2-hours glucose, we suspect the difference in the SNP instruments used may lead to different power and thus different results. Importantly, compared to those previous MR studies [34, 35, 68], our study has the following unique advantages: (**i**) we employed a larger number of valid instruments which were obtained from larger scale GWASs; (**ii**) we performed a more comprehensive analysis by considering a larger set of adult diseases and mediators; (**iii**) we carried out much more extensive sensitivity analyses and simulations to guarantee the robustness of our results, including sensitivity analyses with regard to pleiotropy and maternal effects.

### Public health implications of our results

Our results on the causal effects of birth weight on multiple adult diseases have important implications from a public health perspective. The benefits of reasonably high birth weight in terms of reducing the risks of adult diseases suggest that strategies to increase birth weight can achieve substantial health gains in later life. Indeed, birth weight can be modified via intervening various other modifiable risk factors. For example, cession or reduction in cigarette smoking or drinking during pregnancy can improve birth weight; improved prenatal care and nutrition can improve birth weight; proper supplement of folic acid, iron and multi-mineral vitamins can improve birth weight; educational programs and strict and regular prenatal care can improve birth weight; appropriately longer spacing pregnancies can also decrease the risk of low birth weight [76-79]. Public health policy towards intervening these modifiable risk factors to influence birth weight in the positive direction could reduce disease burden in the adulthood. Importantly, such public health policy towards improving birth weight may have added more benefits in the developing counties than in the developed counties. For example, half of world’s low birth weight infants are born in South Asia [80]; nutrition-based intervention towards improving birth weight [26] there may help curb the unusually high risks of CAD [81], MI [82] and T2D [83] in these developing counties (e.g. India, Pakistan and Nepal). Additionally, as birth weight is often tied with social/economic status [76], some of these strategies intervening the modifiable risk factors to birth weight may have a higher impact in developing countries than in developed countries.

### Limitations of our study

Our analysis results are not without limitations. First, we acknowledge that there was a small overlap between individuals used in the EGG GWAS for birth weight (Table S1) and individuals used in the DIAGRAM GWAS for T2D (Table S7), suggesting that a small set of individuals are simultaneously used to obtain SNP effect size estimates for both birth weight and T2D. In particular, the European Prospective Investigation into Cancer and Nutrition (EPIC) study was included in both these two aforementioned GWASs with an overlapping sample size of ~9,000 individuals (8,939 in EGG and 9,292 in DIAGRAM). Sample overlapping is commonly encountered in GWAS-based MR analysis [84] and can result in model overfitting and biased causal effect estimates. However, the proportion of individuals in the EPIC study is relatively small and represents only 6.22% of the EGG study and 5.86% of the DIAGRAM study, suggesting that the bias resulting from overlapped samples is neglectable [84]. In addition, there is no overlap between samples used in EGG (for birth weight) (Table S1) and samples used in CARDIoGRAMplusC4D (for CAD and MI) (Table S8). Second, for some complex traits, we had to use GWASs with relatively small samples due to data availability reasons. For example, we had to use summary statistics for blood pressures from the Atherosclerosis Risk in Communities (ARIC) GWAS cohort data (Text S2) [55] with only 8,749 individuals. The ARIC sample size is small compared with the previous largest GWAS meta-analysis for blood pressure that includes ~200,000 individuals [85]. However, this largest GWAS for blood pressure only released summary statistics in terms of the absolute effect size estimate but without the effect size direction/sign, and thus cannot be used in the present study. Besides the largest GWAS of blood pressure, we also examined the UK Biobank data [86] and obtained summary statistics available from the online MR-Base platform [87] for blood pressures. Unfortunately, these two data sources contain part of the samples in the EGG study of birth weight without releasing detailed individual overlapping information, and thus cannot be used in the present study. Therefore, we had to use the ARIC data with a relatively small sample size and we emphasize that future research with larger samples to investigate blood pressures will likely be beneficial. Third, like many other MR applications, we have assumed a linear relationship between birth weight and adult diseases. It is certainly possible that non-linear relationships exist; for example, a U-shaped association pattern between birth weight and T2D was observed in a case control study for low birth weight (i.e. birth weight < 2,500g vs. > 2,500g) [13]. However, because birth weights for most individuals collected in the EGG study [37] are in the normal range (95% range is 2,492 - 4,405g; Table S1), a linearity assumption is likely a sensible choice for our study. Fourth, due to the use of GWAS summary statistics, we unfortunately cannot perform stratified analysis by gender and cannot estimate the causal effects of birth weight on adult diseases in males and females separately. Therefore, we are unable to validate different gender specific causal effects of birth weight on adult diseases that are observed in early studies [5]. Fifth, our study focuses only on European population, and future studies are needed to investigate whether our conclusions can be generalized to other human populations.

### Conclusion

Our results suggest that lower birth weight is causally associated with an increased risk of CAD, MI and T2D in later life, supporting the fetal origins of adult diseases hypothesis

## Additional file

Supplementary text for method details, and supplementary Tables and Figures.

## Abbreviations

MR: Mendelian Randomisation
CAD: coronary artery disease
T2D: type 2 diabetes
SNP: single-nucleotide polymorphism
EGG: Early Growth Genetics
GWAS: genome-wide association study
AMD: age-related macular degeneration
PSC: primary sclerosing cholangitis
CKD: chronic kidney disease
CD: Crohn’s disease
UC: ulcerative colitis
PBC: primary biliary cirrhosis
IBD: inflammatory bowel disease
SLE: systemic lupus erythematosus
OR: odds ratio
AS: ankylosing spondylitis
IS: ischaemic stroke
MS: multiple sclerosis
T1D: type 1 diabetes
RA: rheumatoid arthritis
LOO: leave-one-out
MI: myocardial infarction
IVW: inverse-variance weighted
PVE: phenotypic variance explained

## Acknowledgements

We thank the Editor and the reviewers for their thorough and useful comments which have helped to improve our paper. We thank all the GWAS consortium studies for making the summary data publicly available and are grateful of all the investigators and participants contributed to those studies. The GWAS summary data source to these data consortium is given in Text S5 of Supplementary Information.

## Funding

This study was supported by the National Institutes of Health (R01HG009124, R01GM126553), the National Science Foundation (DMS1712933), Youth Foundation of Humanity and Social Science funded by Ministry of Education of China (18YJC910002), the project funded by China Postdoctoral Science Foundation (2018M630607), the Natural Science Foundation of Jiangsu, the Project funded by Postdoctoral Science Foundation of Xuzhou Medical University, QingLan Research Project of Jiangsu for Outstanding Young Teachers, the National Natural Science Foundation of China (81402765), the Statistical Science Research Project from National Bureau of Statistics of China (2014LY112), and the Priority Academic Program Development of Jiangsu Higher Education Institutions (PAPD) for Xuzhou Medical University.

## Authors’ contributions

PZ and XZ conceived the idea for the study. PZ obtained the genetic data. PZ and XZ developed the study methods. PZ performed the data analyses. PZ and XZ interpreted the results of the data analyses. PZ and XZ wrote the manuscript.

## Availability of data and materials

Data sharing not applicable to this article as no datasets were generated or analysed during the current study.

## Ethics approval and consent to participate

Not applicable.

## Consent for publication

Not applicable.

## Competing interests

The authors declare that they have no competing interests.

